# AICAR prevents doxorubicin-induced heart failure in rats by ameliorating cardiac atrophy and improving fatty acid oxidation

**DOI:** 10.1101/2022.08.17.504253

**Authors:** Anurag Choksey, Benjamin D. Thackray, Vicky Ball, Lea Hong Tuan Ha, Eshita Sharma, Brett W. C. Kennedy, Ryan D. Carter, John Broxholme, Michael P Murphy, Lisa C Heather, Damian J Tyler, Kerstin N Timm

**Author notes:** Corresponding author contact: Kerstin Nina Timm, PhD, Department of Pharmacology, University of Oxford, Mansfield Road, Oxford OX1 3QT.

## Abstract

Doxorubicin (DOX) is a widely used chemotherapeutic agent that can cause serious cardiotoxic side effects, leading to functional cardiac decline and ultimately, congestive heart failure (HF). Impaired mitochondrial function and energetics are thought to be key factors driving progression into HF. We have previously shown in a rat model of chronic intravenous DOX-administration that heart failure with reduced ejection fraction correlates with mitochondrial loss and dysfunction. Adenosine monophosphate-dependent kinase (AMPK) is a cellular energy sensor, regulating mitochondrial biogenesis and oxidative metabolism, including fatty acid oxidation. We hypothesized that AMPK activation could restore mitochondrial number and function and therefore be a novel cardioprotective strategy for the prevention of DOX-HF. We therefore set out to assess whether 5-aminoimidazole-4-carboxamide 1-β-D-ribofuranoside (AICAR), an activator of AMPK, could prevent cardiac functional decline in this clinically relevant rat model of DOX-HF. In line with our hypothesis, AICAR improved cardiac systolic function. We show that this could be due to normalisation of substrate supply to the heart, as AICAR prevented DOX-induced dyslipidaemia. AICAR furthermore improved cardiac mitochondrial fatty acid oxidation, despite no increase in mitochondrial number. In addition, we found that AICAR prevented excessive myocardial atrophy, and RNAseq analysis showed that this may be due to normalisation of protein synthesis pathways, which are impaired in DOX-treated rat hearts. Taken together, these results show promise for use of AICAR as a cardioprotective agent in DOX-HF to both preserve cardiac mass and improve cardiac function.

## Introduction

Doxorubicin (DOX) is a chemotherapeutic antibiotic of the anthracycline group first isolated in 1969 from *Streptomyces peucetius*^1^. It is regularly used to treat solid and haematological malignancies, including breast cancer, ovarian cancer, and acute myeloid leukaemia in both adult and paediatric patients^2^. Though efficacious, usage of DOX clinically can be restricted by its serious cardiotox side-effect, which can lead to heart failure (HF) with a mortality rate of 60% following diagnosis^3^. There are currently no treatments available to prevent DOX-HF, and standard heart failure medication is the only option once functional decline is already apparent^4^. Therefore, targeted therapeutics that can prevent DOX-HF are crucial in reducing not only the incidence of DOX-HF but in improving chemotherapeutic regimens and cancer outcomes^5^.

There are a myriad of proposed mechanisms to account for DOX-cardiotoxicity, thought to be distinct from its antitumour inhibition of DNA-synthesis mediated by topoisomerase II inactivation^5^. Indeed, oxidative stress^6^, calcium dysregulation^7^, apoptosis^8^, autophagy^9^ and mitochondrial dysfunction^10^ have all been suggested as mechanisms of DOX-induced cardiotoxicity, with oxidative stress being the predominant theory^11,12^. However, our previous work in a chronic intravenous rat model of DOX-HF, which resembles a clinical schedule of DOX administration seen in patients with cancer, found oxidative stress unlikely to be the main cause of DOX-HF^13^. Our study instead showed a drop in mitochondrial number and function following DOX administration. Therefore, we hypothesised that the mechanism behind DOX-HF may be more substantially related to a loss of mitochondria and decrease in mitochondrial oxidative metabolism^13^, suggesting that reversal of this phenomenon may afford an avenue to abrogate or even treat DOX-HF.

The healthy heart predominantly uses fatty acids for energy generation, accounting for around 70% of total ATP production through oxidative phosphorylation, while glucose, pyruvate and lactate account for less than 30% of cardiac energy utilisation^14^. Importantly, fatty acids are a more efficient energy store than carbohydrates and fatty acid-derived acetyl-CoA inhibits pyruvate dehydrogenase (PDH), thereby limiting glucose oxidation if fatty acids are in plentiful supply. This reciprocal regulation of substrate oxidation is known as the Randle cycle^15^. Activated adenosine monophosphate kinase (AMPK) helps regulate the Randle cycle by stimulating oxidative metabolism and fatty acid uptake^16^. DOX has previously been shown to decrease mitochondrial number and impair cardiac energetics by decreasing levels of activated AMPK^17^. Indeed, decreased AMPK signalling has been proposed as another potential mechanism of DOX cardiotoxicity^18^.

AMPK is a heterotrimeric protein complex, with a catalytic α subunit and regulatory β and γ subunits. AMPK is an important regulator of cellular energy homeostasis as AMP allosterically activates the γ subunit^19^. Further activation of AMPK occurs at phosphorylation sites on both the α and β subunits, through the action of AMPK kinases^20^. AMPK phosphorylation activates peroxisome proliferator-activated receptor-γ coactivator 1α (PGC1α) and PGC1β^21^. Similarly, fatty acid uptake and metabolism is also upregulated both directly and indirectly by AMPK^20^. Therefore, AMPK may provide a mechanistic link between DOX-induced mitochondrial loss and dysfunction, leading to cardiotoxicity and we have previously proposed AMPK activation as a potential mechanism to prevent DOX-HF^23^.

5-aminoimidazole-4-carboxamide-1-β-D-ribofuranoside (AICAR) is a prodrug, which is phosphorylated intracellularly to the active AMP analogue 5-aminoimidazole-4-carboxamide ribonucleotide (ZMP)^24^. ZMP activates AMPK by binding to AMPK’s allosteric site and promoting phosphorylation of its regulatory subunits^24–26^. As an AMPK activator, AICAR has been shown to increase mitochondrial biogenesis^27,28^ and fatty acid oxidation^29,30^, though this has not previously been demonstrated in the context of DOX-HF. Therefore, we set out to determine whether adjuvant AICAR-treatment in rats on DOX-chemotherapy would be sufficient to provide protection against DOX-HF. Here we show that AICAR can ameliorate DOX-HF in rats, potentially by abrogating cardiac atrophy and improving mitochondrial fatty acid oxidation, thereby preserving cardiac systolic function. Taken together, these results show promise for use of AICAR as a cardioprotective agent in DOX-HF.

## Methods

### Animal studies and plasma analysis

All animal experiments conformed to Home Office Guidance on the Operation of the Animals (Scientific Procedures) Act, 1986 and were approved by a local ethics committee. Age and weight-matched male Wistar rats (6-8 weeks, ~250 g starting weight) were treated for 5 consecutive weeks with either saline (n=6) or 3 mg kg^-1^ DOX (n=18). The DOX group was further divided into 2 groups (n=9 each group): one group receiving daily s.c. injections of sterile saline and one group receiving daily s.c. injections of 0.5 g kg^-1^ 5-aminoimidazole-4-carboxamide ribonucleotide (AICAR) starting 48 h before the first dose of DOX and for the full 6 weeks of the study (Figure 1a). Rats were weighed weekly during the study and food intake was measured in one cage per condition (3 rats per cage) for the duration of the 6-week protocol. At weeks 1, 3 and 6 rats were anaesthetized with 2% (v/v) isoflurane with medical oxygen and blood samples were taken for analysis of plasma non-esterified fatty acids (NEFA), triglycerides (TAG) and β-hydroxybutyrate with commercially available kits (Randox, Crumlin, UK). Rats then underwent an MR imaging protocol (see below). At the end of the study rats were sacrificed and their epididymal fat pads weighed and tibia length measured. Some of these rats (n=6 per condition) also underwent mitochondrial isolation and functional analysis (see below).

**Figure 1.**
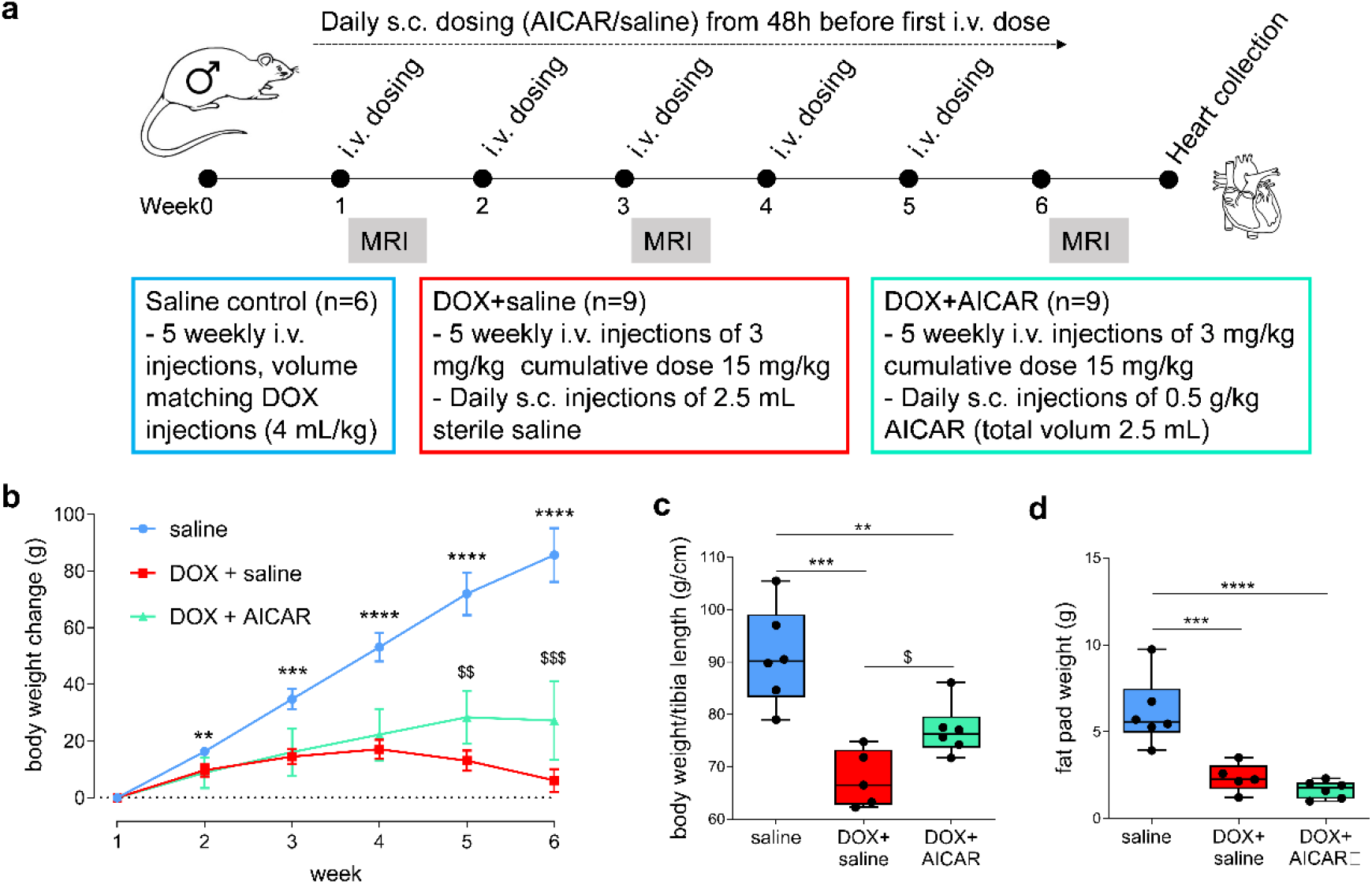
AICAR prevents body weight loss in DOX-treated rats. **a**, Outline of the study. Male Wistar rats were treated intravenously (i.v.) with either 5 weekly injections of saline or 3 mg kg^-1^ doxorubicin (DOX high) and the DOX groups were furthermore split into two groups: one group receiving daily s.c. injections of saline and one receiving daily s.c. injections of 0.5 g kg^-1^ 5-aminoimidazole-4-carboxamide ribonucleotide (AICAR), starting 48 h before the first DOX injection. CINE MR imaging of cardiac function and hyperpolarized [2-^13^C]pyruvate MRS for cardiac metabolic flux measurements were performed at weeks 1, 3 and 6. **b**, Rat body weight change over time normalized to baseline average weight. Data represents mean±SEM. Post mortem analysis of **c**, body weight:tibia length ratio and **d,** epididymal fat pad weight. Box and whisker plots ranging from min to max value with the median indicated by horizontal line. Statistical comparison by one-way ANOVA (**c-d**) with Tukey’s HSD correction method for multiple comparisons. \*\**P*<0.01, \*\*\**P*<0.001, \*\*\*\**P*<0.0001 compared to saline group. ^$^Statistically significant difference between DOX+AICAR and DOX+saline group.

### Cardiac functional and metabolic analysis with MRI and hyperpolarized MRS

After blood sampling, and during the same anaesthesia, rats underwent cardiac functional analysis using CINE MR imaging (MRI) and real-time metabolic flux measurements with hyperpolarized [2-^13^C]pyruvate MR spectroscopy (MRS), performed on a 7 T preclinical MR system (Varian) as previously described^31^. The order of CINE MRI and hyperpolarized [2-^13^C]pyruvate MRS scans was randomized between rats. For the hyperpolarized experiments, 1 mL of 80 mM [2-^13^C]pyruvate was injected into the tail vain over 10 s. ^13^C MR spectra were acquired from the heart every second for 60 s using a 72 mm dual-tuned birdcage volume transmit ^1^H/^13^C coil and a ^13^C two-channel surface receive coil (Rapid Biomedical; 15° hard pulse; 17.6 kHz bandwidth). Multi-coil spectra were combined using an in-house Whitened Single-Value Decomposition script^32^ in Matlab (Mathworks, Natick, MA, USA). The first 30 s of spectra from the first appearance of the pyruvate peak were summed and quantified with AMARES in jMRUI as described previously^33^.

### Mitochondrial analysis

Rats were sacrificed under terminal isoflurane anaesthesia (5% (v/v) in medical oxygen) at week 6 and their hearts excised. A ~100 mg piece of the heart apex was freeze clamped with liquid nitrogen-cooled Wallenberger tongs and from the remaining tissue subsarcolemmal (SSM) and interfibrillar (IFM) mitochondria were isolated^34^. Oxygen consumption rates were assessed in mitochondria (0.15 mg mitochondrial protein per experiment) at 30°C with a Clarke-type oxygen electrode (Strathkelvin Instruments Ltd, Glasgow, UK) using pyruvate+malate (PM), glutamate (G) or palmitoyl-CoA+carnitine (PCC) as substrates. From the frozen apex, 30 mg was used for DNA extraction using a DNeasy^®^blood and tissue extraction kit (Qiagen, Venlo, Netherlands). Quantitative real-time PCR was performed using the Power SYBR Green PCR Master Mix and a Step-one Plus Real-Time PCR system (Thermo Fisher Scientific) to assess copy numbers for the mitochondrial gene cytochrome b (cytB) and the nuclear-encoded gene glyceraldehyde-3-phosphate dehydrogenase (GAPDH). Primer sequences (5’–3’) were as follows: GAPDH sense, AGTATGTCGTG GAGTCTACTGGTG; GAPDH anti-sense, TGAGTTGTCATATTTCTCGTGGTT; cytB sense, GGGTATGTACTCCCATGAGGAC; cytB anti-sense, CCTCCTCAGATTCATTCGAC. The data were analyzed with the comparative C_t_ method^35^ with relative cytB copy number as a marker of mitochondrial number. The remaining heart apex tissue was ground to a fine powder under liquid nitrogen and used for protein and RNA extraction (see Western Blotting and RNAseq) and for citrate synthase activity measurements using standard spectrophotometric analysis techniques^36^.

### Western blotting

Protein was extracted with 100 mg mL^-1^ lysis buffer (75 mM Tris-HCL pH 6.8, 4 M urea, 3.8% SDS, 20% glycerol, 0.1% TritonX-100) containing protease and protein phosphatase inhibitor cocktails (Sigma Aldrich, Gillingham, Dorset, UK). Protein concentrations were measured with BCA protein assay and samples were boiled for 5 min at 90°C after addition of 5%β-mercaptoethanol. A total amount of 15 μg protein was used for SDS-PAGE separation by electrophoresis and proteins were transferred onto nitrocellulose membranes (VWR, Lutterworth, UK). Even protein loading and transfer were confirmed by Revert 700 total protein staining (Li-COR, Cambridge, UK). Primary antibodies (all from rabbit, 1:1000 in 5% BSA): AMPKα (2603), pAMPKα (T172, 2535) (Cell Signalling, Leiden, Netherlands); PGC1α/PGC1β (ab72230). Secondary antibody (goat-anti rabbit HRP 1:10000 in 5% BSA) was purchased from Abcam (Cambridge, UK). Bands were quantified using LICOR C-Digit chemiluminescent detection system (LICOR Biotechnology, Lincoln, Nebraska) and Image Studio software version 5.2.5 (LI-COR).

### RNA extraction and RNAseq library preparation

RNA was extracted with RNeasy^®^ Fibrous Tissue Mini kit (Qiagen, Manchester, UK) using ~30 mg of snap-frozen heart tissue from DOX+saline and DOX+AICAR rats (n=6 each). Material was quantified using RiboGreen (Invitrogen) on the FLUOstar OPTIMA plate reader (BMG Labtech) and the size profile and integrity analysed on the 2200 (Agilent, RNA ScreenTape). RIN estimates for all samples were above 7. Input material was normalised to equal input of 100 ng prior to library preparation. Polyadenylated transcript enrichment and strand specific library preparation was completed using NEBNext Ultra II mRNA kit (NEB) following manufacturer’s instructions. Libraries were amplified on a Tetrad (Bio-Rad) using inhouse unique dual indexing primers (based on DOI: 10.1186/1472-6750-13-104). Individual libraries were normalised using Qubit, and the size profile was analysed on the 2200 or 4200 TapeStation. Individual libraries were normalised and pooled together accordingly. The pooled library was diluted to ~10 nM for storage. The 10 nM library was denatured and further diluted prior to loading on the sequencer. Paired end sequencing was performed using a HiSeq4000 75bp platform (Illumina, HiSeq 3000/4000 PE Cluster Kit and 150 cycle SBS Kit), generating a raw read count of 22 million read pairs per sample.

### RNAseq mapping and counts

RNAseq read pairs were aligned to Rattus norvegicus reference genome, Rnor_6.0 using a splice-aware aligner, Hisat2 version-2.0.4^37^. Gene annotation files were downloaded in GTF format from Ensembl, release 81^38^. Read fragments mapping to annotated exon features were quantified with featureCounts^39^, part of subread-v1.5.0^40^, using default parameters. Values for duplication rates and median 3’ bias were estimated using MarkDuplicates, and CollectRnaSeqMetrics respectively, both implemented in Picard tools v1.92^41^. Normalized read counts and count based metrics were obtained using in-house R scripts^42^.

### RNAseq analysis

We used the above new RNAseq dataset as well as our previously published RNAseq dataset^13^ (Gene Expression Omnibus, accession code GSE154603) for analysis. Raw sequencing count tables were filtered to analyse those with >10 transcripts per million across at least 5 samples. Differentially expressed genes were identified using the DEseq2 R package^43^ using a false discovery rate of 0.05. The gene set was annotated with the org.Rn.eg.db package, and genes were ranked by the test statistic from DESeq2. Gene set enrichment analysis (GSEA) was carried out using the fgsea R package^44^ to identify changes in pathway expression between treatment groups using the C2 Reactome gene set from the Molecular Signatures Database v7.2 (Broad Institute).

### Statistical analysis

Statistical analysis was performed in Prism 6.0 (GraphPad, La Jolla, CA, US). Unpaired Student’s *t*-tests, one-way or two-way ANOVA with Tukey’s HSD adjustment method for multiple comparisons was used and performed as indicated in the figure legends. Significance was assumed at *P* < 0.05. Only significances from multiple comparisons are displayed in the figures.

## Results

### AICAR prevents excessive body weight loss in doxorubicin-treated rats

We first assessed whether daily AICAR injections led to changes in body weight gain or adiposity. We found that while DOX-treated rats that only received subcutaneous injections of saline lost weight after the 4^th^ injection of DOX, this was prevented in the DOX+AICAR group (Figure 1b). At the final timepoint, the body weight to tibia length ratio was significantly lower in both the DOX+saline and DOX+AICAR group compared to the saline control group, however, there was a significant increase in body weight to tibia length ratio in the DOX+AICAR group compared to the DOX+saline group (Figure 1c). This may be due to improved appetite/reduced nausea as we found that DOX+saline rats consumed less food during the last two weeks of the study, while DOX+AICAR rats maintained food intake throughout the 6 weeks (Figure S1). However, rats treated with DOX+AICAR consumed overall less food (5.85 ± 0.44 g 100 g^-1^ rat day) compared to rats treated with DOX+saline (6.32 ± 1.66 g 100 g rat^-1^ day), with saline control rats consuming the most (7.25 ± 0.64 g 100 g rat^-1^ day). We furthermore found that the epididymal fat pad weight was significantly reduced in both the DOX+saline and DOX+AICAR group compared to the saline control group, and there was no difference between the DOX+saline and DOX+AICAR group (Figure 1d). AICAR thus prevents overall body weight loss despite a reduction in overall food intake normalised to body weight, but does not increase fat stores. This suggests less loss of lean body mass in the DOX+AICAR group than was seen in DOX+saline rats.

### AICAR ameliorates cardiac dysfunction and atrophy in DOX-HF

We next assessed whether AICAR could also improve cardiac function by performing CINE MRI at weeks 1, 3 and 6 of the protocol. Cardiac end-systolic volume was increased in DOX+saline treated rats at week 6 but this was not the case in the DOX+AICAR rats (Figure 2a). End-diastolic volume was decreased in both DOX+saline and DOX+AICAR rats at weeks 3 and 6 (Figure 2b) compared to controls. Cardiac left-ventricular ejection fraction (LVEF) was preserved in AICAR-treated DOX rats between weeks 3 and 6, while DOX+saline rats had a significant decrease in LVEF at weeks 3 and 6 (Figure 2c). Stroke volume was decreased at 3 and 6 weeks in both the DOX+saline group and DOX+AICAR groups vs. healthy controls, however the reduction in stroke volume of the DOX+AICAR group was significantly improved compared to DOX+saline animals at week 6 (Figure 2d). Heart rate was decreased in DOX+AICAR rats at weeks 3 and 6, but only at week 6 in DOX+saline rats (Figure 2e). Cardiac output was decreased in both DOX+saline and DOX+AICAR rats at weeks 3 and 6, however, cardiac output was significantly higher in DOX+AICAR rats at week 6 compared to DOX+saline rats (Figure 2f). These data suggest that AICAR confers cardioprotection in DOX-treated rats by improving systolic function. Interestingly, while both DOX+AICAR and DOX+saline rats displayed a decreased heart weight:tibia length ratio, indicative of cardiac atrophy at week 6, there was a significant increase in the ratio of DOX+AICAR rats compared to DOX+saline rats (Figure 2g), suggesting some preservation in cardiac mass. Indeed, upon interrogating our previous RNAseq dataset^13^ (Gene Expression Omnibus, accession code GSE154603) we found that DOX-hearts showed upregulation of pathways relating to rRNA processing, protein elongation and protein targeting, which could indicate impaired ribosomal function (Figure S2a). This could result in reduced protein synthesis, suggesting a mechanism by which DOX causes cardiac atrophy. In comparison to DOX+saline rats, these pathways were downregulated by DOX+AICAR (Figure S2b), indicating a potential rescue effect on protein metabolism, and explaining preservation of myocardial mass due to normalisation of protein synthesis pathways. DOX+AICAR rats furthermore showed increased cardiac expression of pathways relating to extracellular matrix organisation, protein glycosylation and transport to the Golgi apparatus, which may indicate alterations in protein processing and cardiac remodelling. Taken together, we hypothesised that these results suggest AICAR is mildly cardioprotective through conservation of myocardial mass in DOX-treated rats, and that this may be due to AMPK activation leading to increased mitochondrial number, improved substrate oxidation (Figure 2h), and amelioration of cardiac atrophy by restoration of energy supply and normalised protein synthesis.

**Figure 2.**
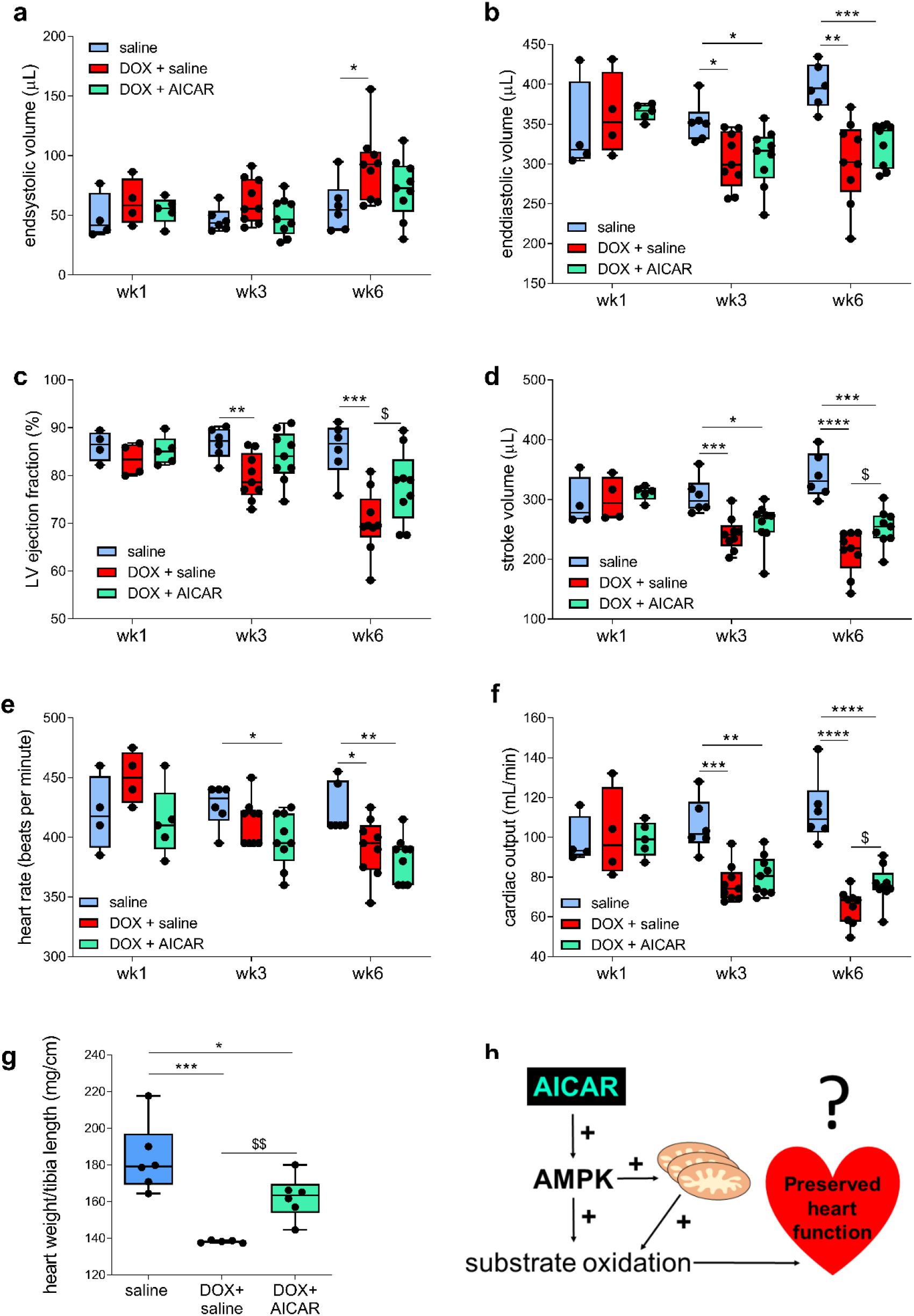
AICAR improves cardiac function and mass in DOX-treated rats. CINE MRI in rats at weeks 1, 3 and 6. Left ventricular (LV) **a**, end-systolic volume **b**, end-diastolic volume **c,** LV ejection fraction **d,** stroke volume **e,** heart rate **f**, cardiac output **g,** heart weight:tibia ratio and in rats from all three groups. **h,** Cartoon depicting working hypothesis that AICAR preserves cardiac function either by increasing mitochondrial number or by increasing substrate oxidation or a combination of the two, after AMPK activation. Box and whisker plots ranging from min to max value with the median indicated by horizontal line. Statistical comparison by one-way ANOVA (**g-h**) or two-way ANOVA (**a-f**) with Tukey’s HSD correction method for multiple comparisons. \**P*<0.05, \*\**P*<0.01, \*\*\**P*<0.001, \*\*\*\**P*<0.0001 compared to saline group. ^$^Statistically significant difference between DOX+AICAR and DOX+saline group.

### AICAR does not improve myocardial carbohydrate metabolism or mitochondrial number

To explore the mechanism of AICAR-mediated cardioprotection, we assessed activation of AMPK and downstream targets by Western Blotting (Figure 3a-e). To our surprise, we found no increase in cardiac AMPK phosphorylation with AICAR, and no difference in expression of AMPK (Figure 3b-c). We also assessed targets downstream of AMPK activation, PGC1α and PGC1β, and found that whilst there was no difference in PGC1α between groups, DOX+saline and DOX+AICAR rats showed a decrease in myocardial PGC1β expression (Figure 3d-e). As PGC1β upregulates mitochondrial biogenesis, its repression in DOX+AICAR rats suggests that AICAR does not act cardioprotectively by restoring mitochondrial number in DOX+AICAR hearts. In tandem with this observation, mitochondrial number was not improved in DOX+AICAR hearts vs. DOX+saline hearts, when assessing a marker of mitochondrial DNA (cytochrome b) compared to a nuclear DNA marker (GAPDH), or when using citrate synthase activity as a proxy for mitochondrial number (Figure 3f-g). In concert with this lack of increased mitochondrial number by AICAR, we found that injection of hyperpolarized [2-^13^C]pyruvate showed a decrease in ^13^C-glutamate labelling in the heart in both DOX+saline and DOX+AICAR rats at week 6, with a statistically significant decrease in glutamate labelling between the DOX+AICAR and DOX+saline groups already apparent at week 3 (Figure 3h-i). We have previously shown that reduced glutamate-labelling from hyperpolarized [2-^13^C]pyruvate as a proxy for TCA-cycle flux is a surrogate *in vivo* imaging marker of mitochondrial number, which is reduced at week 6 of the treatment protocol^13^. T o our surprise, we furthermore found that the cardiac acetylcarnitine:pyruvate ratios were decreased early in the timecourse at weeks 1 and 3 in the DOX+AICAR group compared to the DOX+saline and saline control groups (Figure 3j). Hyperpolarised flux of ^13^C-label from [2-^13^C]pyruvate into acetylcarnitine measures acetyl-CoA buffering capacity via the carnitine pool^45^, and carnitine is important for the mitochondrial import of long-chain fatty acids. Cardiac carbohydrate oxidation is directly affected by fatty acid oxidation via the Randle cycle^15^, and a decrease in the flux from hyperpolarised [2-^13^C]pyruvate into acetylcarnitine could imply increased utilisation of fatty acids in DOX+AICAR rats compared to DOX+saline treated animals. Taken together, these results suggest that AICAR exerts effects on cardiac metabolism that are irrespective of mitochondrial number. Indeed, our previous RNAseq dataset^13^ shows that DOX causes decreases in myocardial mitochondrial biogenesis, TCA cycle and electron transport chain as well as fatty acid metabolism (Figure S2a). To evaluate further the causative role of lipid handling in DOX-HF, we set out to establish whether AICAR did indeed induce alterations in fatty acid oxidation and lipid handling which could explain improvements in cardiac function.

**Figure 3.**
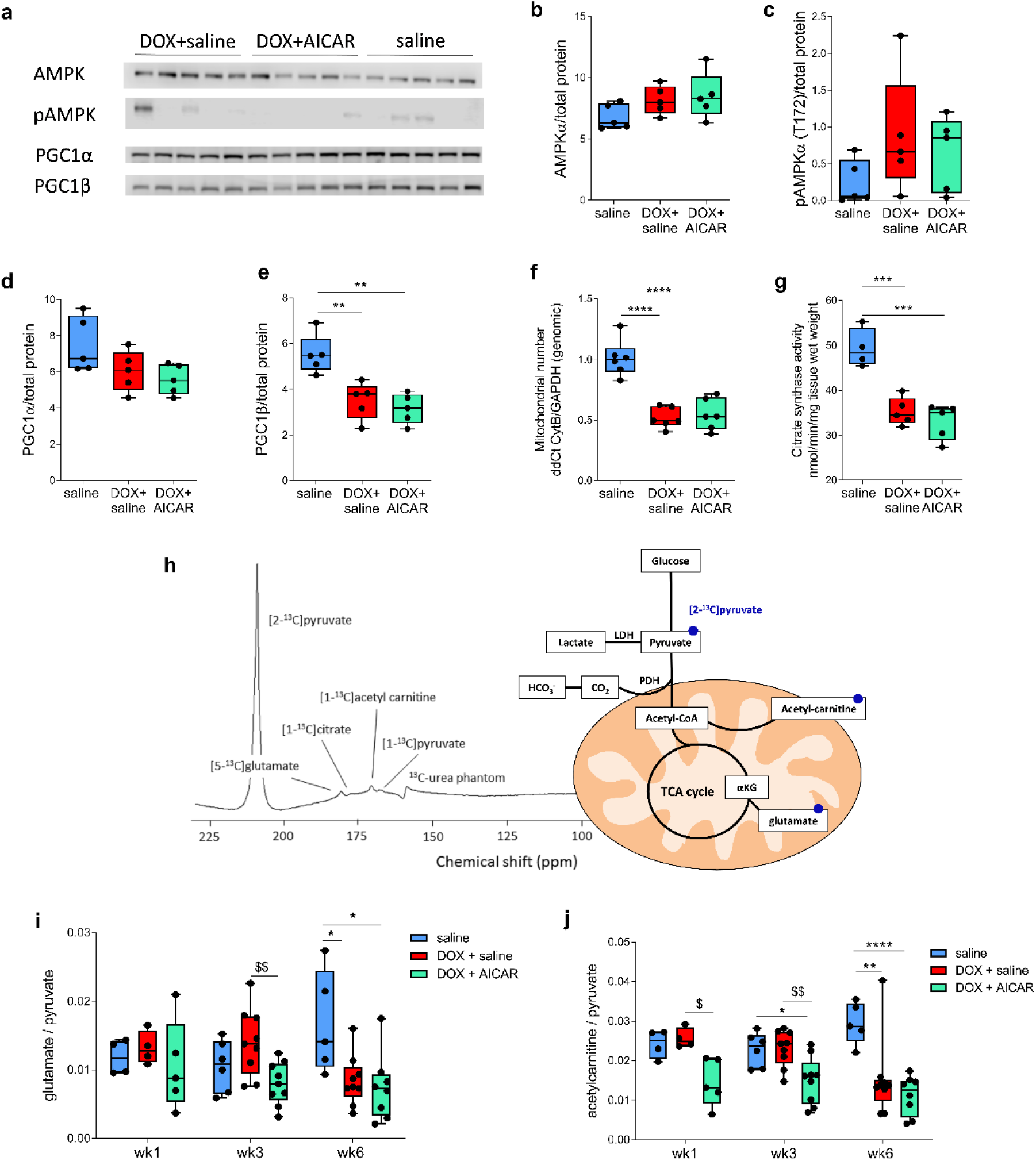
AICAR does not increase oxidative carbohydrate metabolism, AMPK phosphorylation or mitochondrial number in DOX-treated rats. **a**, Western blots of cardiac protein extracts. Bands were normalized to total protein content (Revert total protein stain). **b-g,** Average protein expression of 5 separate protein extracts per treatment condition normalized to total protein for **b**, adenosine monophosphate-dependent kinase (AMPK) **c,** pAMPK **d,** peroxisome proliferator-activated receptor γ coactivator 1α (PGC1α) and **e**, PGC1β. **f,** Mitochondrial number assessed by qPCR analysis of a mitochondrial gene (cytochrome B; cytB) compared to a nuclear gene (glyceraldehyde-3-phosphate dehydrogenase; GAPDH) in genomic DNA-extracts from heart tissue. **g**, mitochondrial number assessed by citrate synthase enzyme activity in tissue extracts. **h**, Sum of the first 30 s of MR spectra after intravenous injection of hyperpolarized [2-^13^C]pyruvate in male Wistar rats treated for 5 consecutive weeks with intravenous weekly injections of saline or 3 mg kg^-1^ DOX (DOX high) and with daily s.c. injections of either saline or 0.5 g/kg AICAR. **i**, Cardiac glutamate:pyruvate ratio and **j**, acetyl-carnitine:pyruvate ratio of. Box and whisker plots ranging from min to max value with the median indicated by horizontal line. Statistical comparison by one-way ANOVA (**b-g**) or two-way ANOVA (**i-j**) with Tukey’s HSD correction method for multiple comparisons. \**P*<0.05, \*\**P*<0.01, \*\*\**P*<0.001, \*\*\*\**P*<0.0001 compared to saline group. ^$^Statistically significant difference between DOX+AICAR and DOX+saline group.

### AICAR improves lipid handling and myocardial mitochondrial fatty acid oxidation

We have previously reported an increase in plasma non-esterified fatty acid and β-hydroxybutyrate in this rat model in DOX-treated animals, which correlates with impaired cardiac function^13^. We therefore wanted to explore whether rescue of cardiac function by AICAR abrogated these changes. Indeed, while we found an increase in triglyceride (TAG), NEFA and β-hydroxybutyrate in the DOX+saline compared to healthy saline control group, we saw no difference in the DOX+AICAR compared to the saline control groups. There was however a significant decrease of NEFA and β-hydroxybutyrate between the DOX+AICAR and DOX+saline groups, suggesting normalisation of this part of the DOX-HF induced phenotype (Figure 4a-c). Therefore, we next explored whether substrate oxidation in isolated cardiac mitochondria was different between DOX+saline and DOX+AICAR rat hearts. Mitochondrial oxygen consumption measurements showed that AICAR did not prevent reduced carbohydrate and amino acid oxidation in isolated subsarcolemmal mitochondria (SSMs) and interfibrillar mitochondria (IFMs). However, AICAR did normalize fatty acid oxidation in IFMs compared to DOX+saline rats, and was comparable to saline control rats in SSMs, despite no statistically significant increase between the DOX+saline and DOX+AICAR group (Figure 4d-e). Taken together, these results suggest that normalised lipid handling and increased myocardial fatty acid oxidation in DOX+AICAR rats could lead to improved cardiac energetics sustaining cardiac function in this model (Figure 4f).

**Figure 4.**
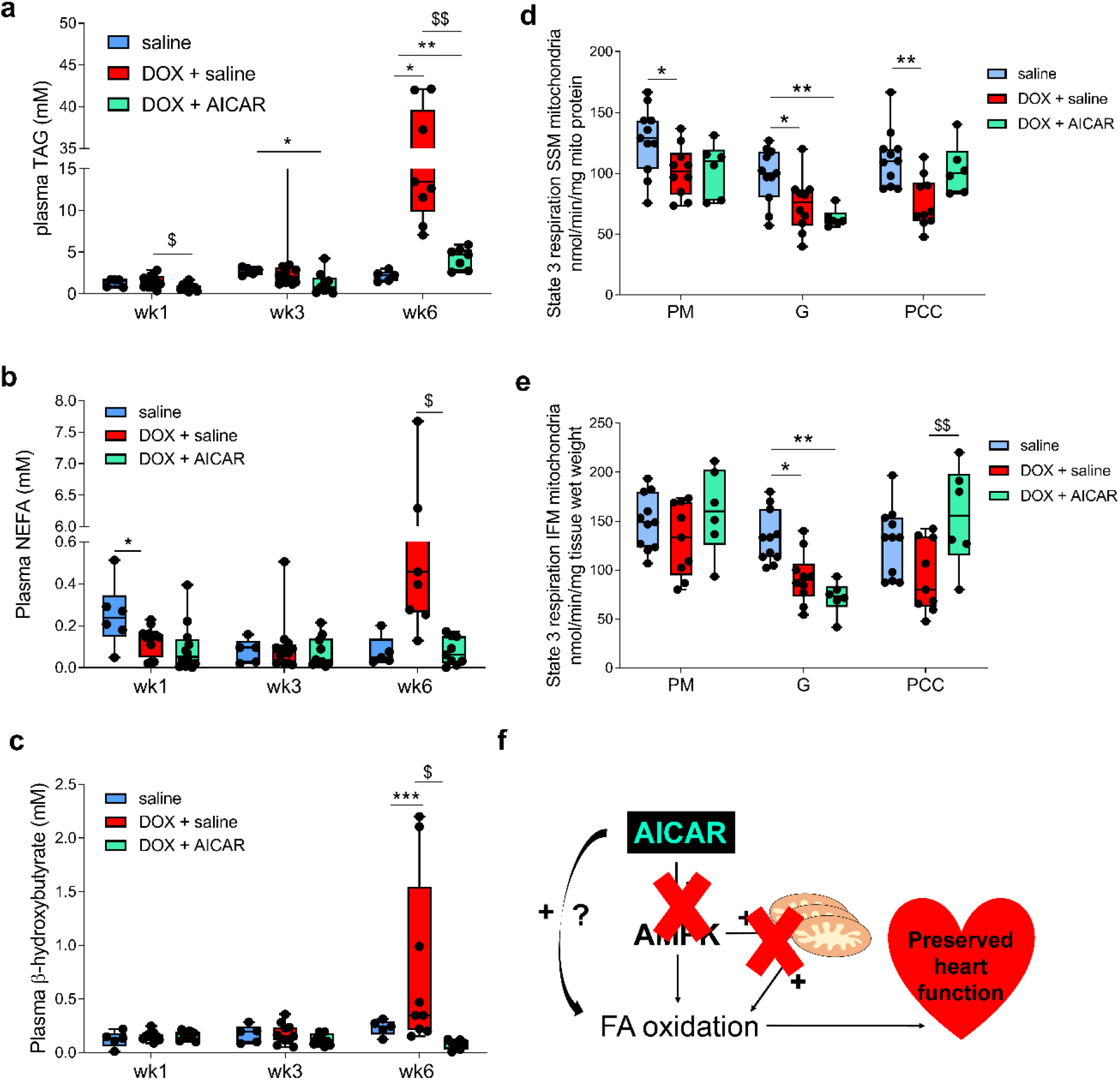
AICAR prevents dyslipidemia and improves mitochondrial fatty acid oxidation in DOX-treated rats. Plasma **a**, triacylglycerides (TAG) **b,** non-esterified fatty acid (NEFA) and **c,** β-hydroxybutyrate levels in rats treated for 5 consecutive weeks with i.v. injections of either saline or 3 mg kg^-1^DOX and the DOX group further treated with either daily s.c. injections of saline or 0.5 g kg^-1^ AICAR. Oxygen consumption measurements (Clarke-style electrode) performed in isolated sub-sarcolemmal (SSMs) and interfibrillar (IFMs) isolated at week 6 (**d-e**). State 3 respiration with pyruvate+malate (PM), glutamate (G) or palmitoyl-CoA + carnitine (PCC) as a fuel in SSMs (**d**) and IFMs (**e**). **f,** cartoon depicting hypothesis of AICAR-cardioprotection. AICAR does not appear to activate AMPK or increase mitochondrial number but improves lipid handing and mitochondrial fatty acid oxidation in DOX-treated rats. Box and whisker plots ranging from min to max value with the mean indicated by horizontal line. Statistical comparison by two-way ANOVA with Tukey’s HSD correction method for multiple comparisons. \**P*<0.05, \*\**P*<0.01, \*\*\**P*<0.001 compared to saline control group. ^$^Statistically significant difference between DOX+AICAR and DOX+saline group.

## Discussion

We show here that AICAR treatment provides partial cardioprotection in a clinically-relevant rat model of DOX-cardiotoxicity. We found that this may be due to improved lipid handing and increased mitochondrial fatty acid oxidation as well as prevention of cardiac atrophy by normalisation of protein synthesis pathways rather than increased mitochondrial biogenesis. Surprisingly, we did not observe AMPK activation by phosphorylation in cardiac tissue. However, AICAR has a short plasma half-life^46^ (15 min in rats) and tissues were collected several hours after the last AICAR injection, making it possible that we may have missed a transient AMPK activation by phosphorylation. Similarly, AMP itself allosterically renders AMPK more active, and phosphorylation only acts additively^47^. AICAR may therefore have an effect on AMPK activity in this model, despite absence of observable changes in phosphorylation state.

We have previously shown with hyperpolarized MRS that TCA cycle flux and acetyl-carnitine buffering capacity is reduced in DOX-treated rats, assessed by ^13^C label incorporation into glutamate and acetylcarnitine from hyperpolarized [2-^13^C]pyruvate, respectively^13^. We concluded that this was due to a decrease in mitochondrial number. In this study however, we could neither see an improvement in PGC1α or β protein levels, nor mitochondrial number in DOX+ACAIR hearts, suggesting that AICAR does not act cardioprotectively by restoring mitochondrial number. Therefore, if the measurements with hyperpolarized [2-^13^C]pyruvate evaluated only mitochondria number, AICAR treatment would be expected to show similar decreases in glutamate and acetylcarnitine labelling in DOX rats. However, we show here that while DOX+AICAR rats displayed a similar effect on glutamate labelling, acetyl-carnitine labelling was significantly reduced early in the treatment timecourse (at weeks 1 and 3), not just at week 6. Taken together, these observations suggest that AICAR causes a metabolic switch in the mitochondria of DOX treated animals, perhaps driving increased oxidation of fatty acids. These changes in substrate utilisation and energy generation may increase mitochondrial energetic efficiency, ameliorating the phenotypic changes seen in DOX-HF without increasing overall mitochondrial number. Indeed, we found that AICAR increased fatty acid oxidation in isolated cardiac mitochondria.

Caloric intake is particularly important to consider in this regard, as decreased nutritional intake will limit the metabolic landscape with which the heart can generate energy. Some of our observations may be explained in part by DOX+AICAR rats maintaining food intake throughout the study (Figure S1), whereas in DOX+saline food intake drops at weeks 5 and 6. However, as the DOX+AICAR group ate less food than the DOX+saline for the first 4 weeks of the study, and ate less food over the 6 week period overall, it is unlikely to fully explain the mechanism of AICAR’s preservation of cardiac function and of plasma lipid content and myocardial fatty acid oxidation. Prolonged AMPK activation improves import of fatty acids through the mitochondrial carnitine transporter^30^, which may explain improved fatty acid oxidation in our model, providing transient AMPK activation was achieved following each AICAR administration over the 6-week period.

However, AICAR also has AMPK-independent functions, namely reducing blood pressure^48^ and increasing the extracellular adenosine pool^49^, which could improve cardiac function by decreasing afterload and increasing inotropy, respectively. Indeed, previous studies on AICAR in the context of myocardial infarction have suggested it maintains contractile function through an AMPK independent, adenosine mediated mechanism^50,51^. AICAR was thought to increase endogenous adenosine levels (though the exact mechanism remains unclear) in a site-specific manner, increasing blood flow to ischaemic tissue and inhibiting immune mediated inflammation^52^. Though we did not assess cardiac adenosine levels in this study, it appears unlikely to be fully responsible for the longer term cardioprotective effect we have seen. Importantly, we found AICAR treatment significantly increased cardiac mass/tibia length ratio. This suggests that longer term cardioprotection is likely related to a maintenance of cardiac mass through AICAR’s intracellular AMPK mediated effects, rather than a purely haemodynamic phenomenon caused by changes in adenosine levels. As AICAR’s effects on the adenosine pool are relatively short lived due to the short half-life of the drug^46^, future analysis of cardiac function after cessation of DOX and AICAR could separate these effects, alongside more direct measurements of intracellular adenosine.

Interestingly, we also found DOX treatment upregulates the expression of pathways relating to protein synthesis and processing, which may initially appear paradoxical as we also saw cardiac atrophy and body weight loss. However, increased expression of genes in these pathways may be maladaptive, leading to cardiac atrophy. Under this line of reasoning, AICAR’s relative downregulation upon the same pathways warrants further investigation as a potential mechanism of cardioprotection by restoring protein synthesis pathways and thus abrogating cardiac atrophy. However, it remains unclear whether any of the changes seen in the RNAseq data are cause-or-effect, as they are taken from whole heart-RNA extracts post-sacrifice and after the onset of DOX-HF. Further, whether these transcriptomic changes translate into changes in functional protein expression remains unclear^53^. This is an important limitation of the data, especially because cell-culture studies have shown DOX to affect the ubiquitination and subcellular localisation of ribosomal proteins, which may play an important role in their function. Overall, this represents an intriguing potential avenue for future study.

## Conclusion

We show here that AICAR can ameliorate cardiac dysfunction and atrophy in a clinically-relevant rat model of DOX-HF. This could be due to prevention of dyslipidaemia and an increase in cardiac fatty acid oxidation capacity, leading to a reciprocal decrease in carbohydrate oxidation and improved cardiac energetics rather than by promoting mitochondrial biogenesis. We also found evidence that ribosomal function and protein synthesis are impaired with DOX, and this may be restored with AICAR, ameliorating atrophy. AICAR has been safely used in patients^54^, making its use as a cardioprotective agent for patients receiving DOX chemotherapy feasible^55^.

## Acknowledgements

K.N.T. would like to acknowledge the Oxford British Heart Foundation Centre of Research Excellence (grant code RE/18/3/34214) for funding this project. L.C.H. would like to acknowledge the British Heart Foundation for funding (grant code FS/17/58/33072). We thank the Oxford Genomics Centre at the Wellcome Trust Centre for Human Genetics (funded by Wellcome Trust grant reference 203141/Z/16/Z with additional support from the NIHR Oxford BRC) for the generation and initial processing of the sequencing data.

## Author contributions

K.N.T. conceptualized the study, designed the experiments and wrote the manuscript with input from M.P.M., L.C.H. and D.J.T.. K.N.T. performed experiments with assistance as follows: V.B. assisted with all MRI experiments and plasma analysis; B.D.T. performed mitochondrial measurements; B.W.C.K. performed enzymatic assays and edited the manuscript; E.S. and J.B. performed initial RNAseq mapping and counts. A.C., L.H.T.H. and R.D.C. performed RNAseq analysis. A.C. furthermore performed Western blotting and co-wrote the manuscript.

