## Supplementary figures for "AICAR prevents doxorubicin-induced heart failure in rats by ameliorating cardiac atrophy and improving fatty acid oxidation"

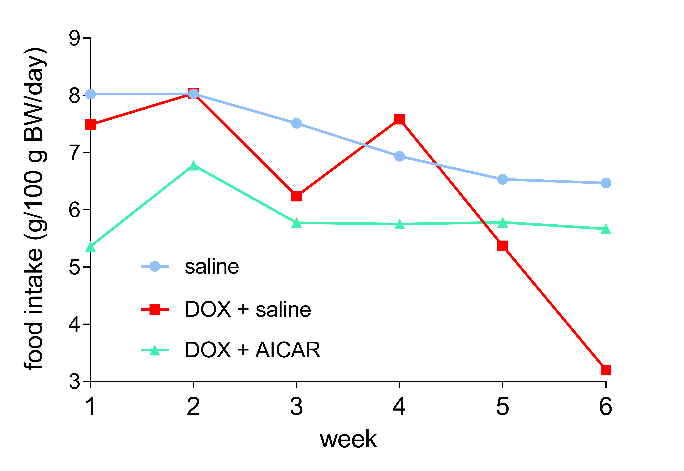

**Figure S1. Food intake in rats.**

Food intake assessed per rat cage and calculated as g food eaten/100g rat/day.

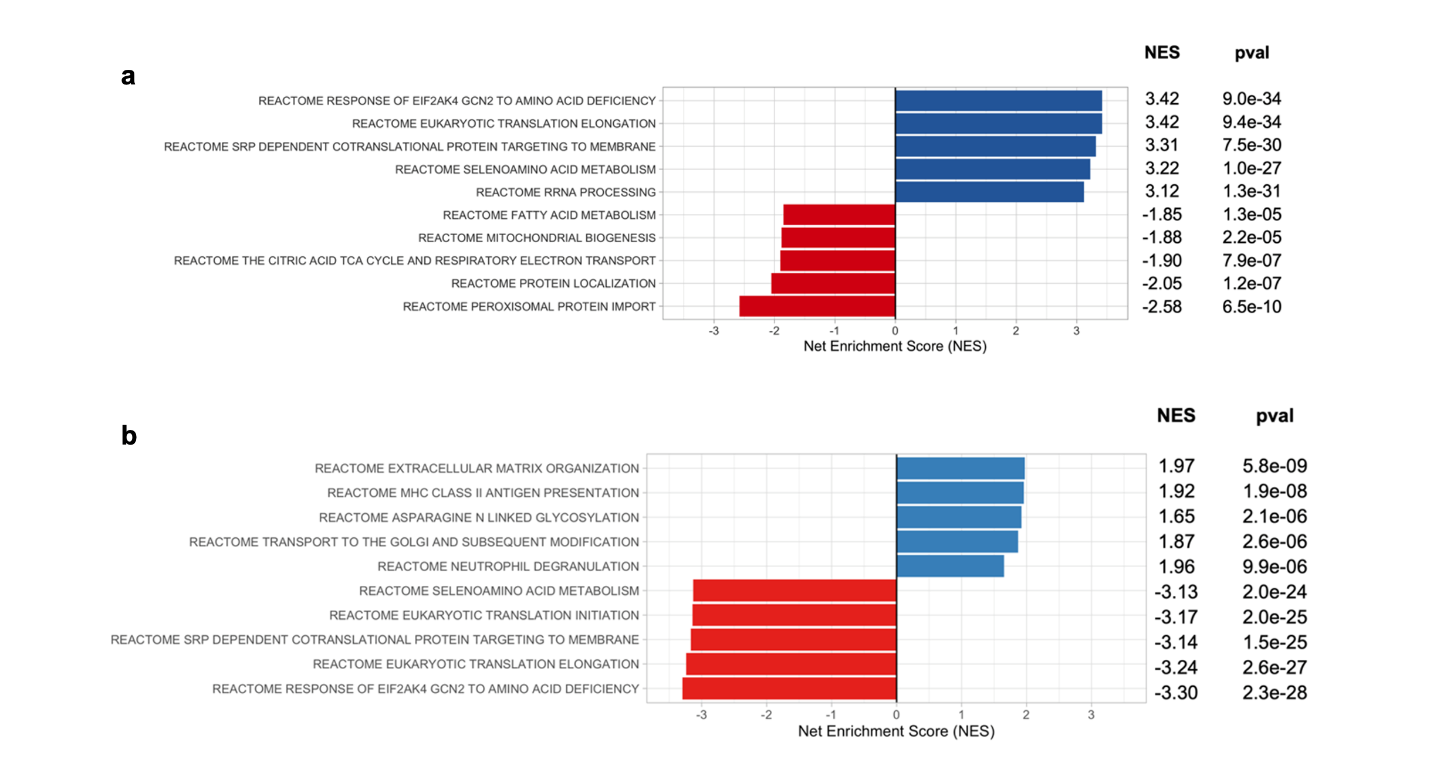

**Figure S2. Protein synthesis pathways are upregulated in DOX treated rats and restored by AICAR.** GSEA results of (a) DOX-treated rats compared to saline controls (dataset on Gene Expression Omnibus, accession code GSE154603)^17^ and (b) DOX+AICAR treated rats compared to DOX-treated rats, using the C2 Reactome gene set five most up- (blue) and down-(red) regulated pathways (determined by p-value), ordered by net enrichment score (NES).
